# The ACE2-binding interface of SARS-CoV-2 Spike inherently deflects immune recognition

**DOI:** 10.1101/2020.11.03.365270

**Authors:** Takamitsu Hattori, Akiko Koide, Tatyana Panchenko, Larizbeth A. Romero, Kai Wen Teng, Takuya Tada, Nathaniel R. Landau, Shohei Koide

## Abstract

The COVID-19 pandemic remains a global threat, and host immunity remains the main mechanism of protection against the disease. The spike protein on the surface of SARS-CoV-2 is a major antigen and its engagement with human ACE2 receptor plays an essential role in viral entry into host cells. Consequently, antibodies targeting the ACE2-interacting surface (ACE2IS) located in the receptor-binding domain (RBD) of the spike protein can neutralize the virus. However, the understanding of immune responses to SARS-CoV-2 is still limited, and it is unclear how the virus protects this surface from recognition by antibodies. Here, we designed an RBD mutant that disrupts the ACE2IS and used it to characterize the prevalence of antibodies directed to the ACE2IS from convalescent sera of 94 COVID19-positive patients. We found that only a small fraction of RBD-binding antibodies targeted the ACE2IS. To assess the immunogenicity of different parts of the spike protein, we performed *in vitro* antibody selection for the spike and the RBD proteins using both unbiased and biased selection strategies. Intriguingly, unbiased selection yielded antibodies that predominantly targeted regions outside the ACE2IS, whereas ACE2IS-binding antibodies were readily identified from biased selection designed to enrich such antibodies. Furthermore, antibodies from an unbiased selection using the RBD preferentially bound to the surfaces that are inaccessible in the context of whole spike protein. These results suggest that the ACE2IS has evolved less immunogenic than the other regions of the spike protein, which has important implications in the development of vaccines against SARS-CoV-2.

## Introduction

The severe acute respiratory syndrome coronavirus 2 (SARS-CoV-2) has caused a worldwide outbreak of the coronavirus disease 2019, COVID-19. As of August 1st 2020, over 17 million cases have been confirmed globally, leading to 675,060 deaths (https://www.who.int/emergencies/diseases/novel-coronavirus-2019/situation-reports/). A number of drugs and vaccines for COVID-19 are currently in clinical trials, yet no agents or vaccines have been approved by the government agencies. As such, the host immune responses remain the main source of protection against the virus. Still many aspects of the immune responses and the strategies employed by the virus to evade them are unknown.

The entry of SARS-CoV-2 into host cells is mediated by a virus-surface spike glycoprotein that forms a trimer (Fig.1A) [1, 2]. The spike protein is composed of two subunits, S1 and S2. The S1 subunit interacts with the host cell receptor, angiotensin-converting enzyme 2 (ACE2), a crucial step in cell attachment, whereas the S2 subunit plays a role in the fusion of the viral and cellular membrane, a crucial step in cell entry [2, 3]. The receptor binding domain (RBD) in the S1 subunit is the domain that binds ACE2 with an affinity in the low nanomolar range [2, 4, 5], and this interaction initiates the conformational change of the spike protein from the pre-fusion state to the post-fusion state. Consequently, antibodies targeting the ACE2-interacting surface (ACE2IS) located in the RBD of the spike protein can compete with the RBD-ACE2 interaction, serving as neutralizing antibodies. Indeed, such antibodies isolated from COVID-19 patients showed potent neutralization effects and are being developed as therapeutics [6–9]. Thus, eliciting antibodies targeting the ACE2IS from the immune system should be critical for controlling and preventing COVID-19.

**Figure 1.**
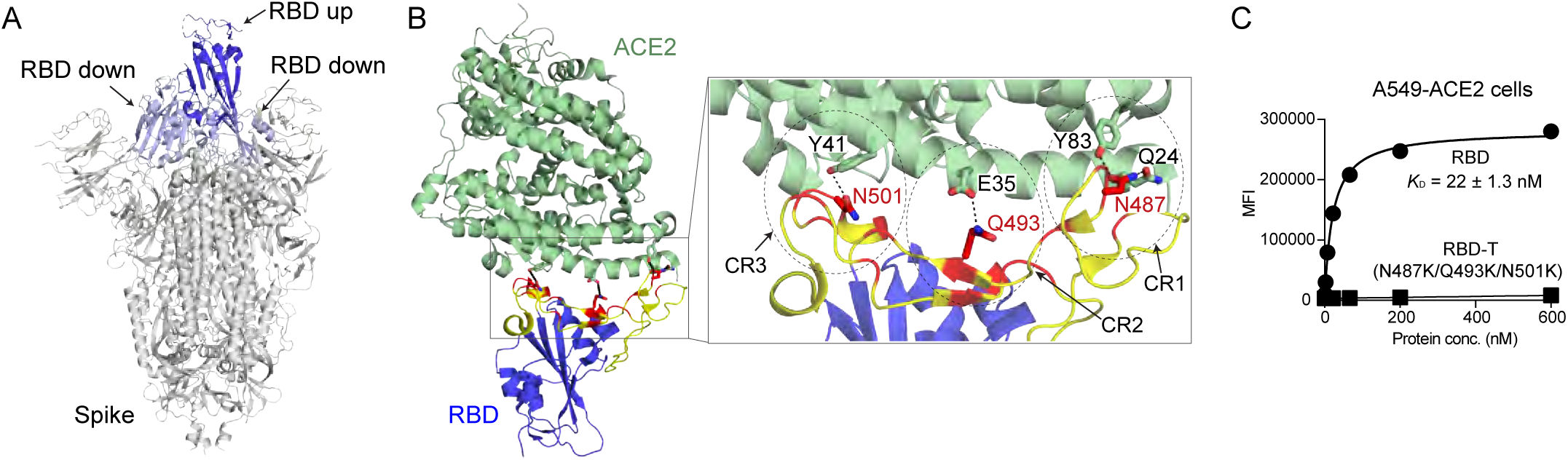
Design of an RBD triple mutant that disrupts the ACE2IS. (A) The structure of the spike trimer (PDB 6VSB). The RBD in the up and down conformations are shown in blue and light blue, respectively. (B) The RBD in complex with ACE2 (PDB 6M0J). The RBD core is shown in blue and ACE2 is shown in green. The receptor binding motif in the RBD is shown in yellow, and the residues contacting ACE2 are shown in red. The dotted circles indicate contact regions (CR1, CR2 and CR3). The amino acid residues (N487, Q493 and N501) in the RBD are shown as stick model and labeled in red, and their contacting residues in ACE2 are labeled in black. (C) Binding titration of the RBD and RBD-T to ACE2 expressing A549 cells. Data shown here are from triplicate measurements. The *K*D values are from curve fitting of a 1:1 binding model, and the errors shown are the s.d. from curve fitting. Error bars are within the size of the symbols.

A region of an antigen is considered immunogenic if it is readily recognized by antibodies, although many aspects in the immune responses of the host can influence the overall immunogenicity. From the viewpoint of viral evolution, there should be a strong selective pressure favoring the protection of a region that is crucial for the viral life-cycle from host immune attacks. For example, the CD4-binding site of the HIV gp120 envelope glycoprotein is situated in a deep cleft heavily protected by glycosylation [10]. However, studies revealed that the spike proteins of coronavirus including SARS-CoV-2, SARS-CoV and MERS-CoV are not densely glycosylated, unlike HIV-1 envelope, Lassa virus glycoprotein complex, and influenza hemagglutinin envelop [11, 12]. In particular, the ACE2IS in the SARS-CoV-2 spike protein appears not highly protected with glycosylation [12]. Intriguingly, a number of studies of SARS-CoV-2 specific antibody responses in COVID-19 patients have revealed that, although most convalescent plasmas from COVID-19 patients contain antibodies targeting the spike protein and the RBD, they have low neutralization activity [13–15]. Thus, analysis of the immunogenic surfaces, or the “immunogenicity landscape”, of the spike protein and the RBD in terms of eliciting high-affinity antibodies should provide important insights into the antibody response to COVID-19 as well as vaccine design, because the spike protein and the RBD are the antigens in most first-generation vaccines for COVID-19 (https://www.who.int/publications/m/item/draft-landscape-of-covid-19-candidate-vaccines).

In this study, to define immunogenic properties of epitopes within the spike protein, we performed *in vitro* selection of a synthetic human antibody library against the spike protein and the RBD. We designed an RBD mutant that abolished binding of the RBD to ACE2, and utilized it as a tool for epitope analysis of convalescent sera and in antibody discovery. Our results suggest that the ACE2IS in the RBD is a minimally immunogenic surface within the spike protein, and we discuss the molecular underpinning of this finding and its implications for vaccine design.

## Results and Discussion

### Design of an RBD mutant that disrupts the ACE2IS

To develop a tool for facilitating the analysis of ACE2IS-binding antibodies, we designed an RBD mutant that disrupts the ACE2IS. The ACE2IS in the RBD is mainly composed of three contact regions, CR1, CR2 and CR3 (Fig.1B) [16]. From each contact region, we chose a key residue (N487, Q493 and N501) that forms hydrogen bond(s) with residues in ACE2, and mutated them to Lys residues to disrupt both electrostatic and van der Waals interactions (Fig. 1B, Supplementary Fig.1). We term this mutant as RBD-T hereafter. Binding analysis using ACE2-expressing cells showed that the RBD bound to ACE2 with high affinity (*K*_D_ ∼ 20nM), consistent with previous reports [1, 2, 5], and that RBD-T nearly completely lost binding to ACE2 (Fig. 1C). Likewise, these mutations should disrupt antibody binding to the ACE2IS. A binding assay using the RBD and RBD-T is a straightforward method to determine whether antibodies recognize the ACE2IS or not, and thus we utilized the RBD-T as an epitope analysis tool for anti-RBD antibodies.

### Analysis of convalescent serum using RBD-T

To characterize the prevalence of antibody response to the ACE2IS in COVID-19 patients, we performed binding analysis of 94 convalescent serum samples [14] from healthcare workers who showed PCR-positivity for COVID-19. In this experiment, we measured the IgG binding to the RBD in the absence and presence of excess soluble RBD-T competitor.

Strikingly, more than 60% of the signal was lost in the presence of the RBD-T competitor in 80% of the serum samples, indicating that the majority of the antibodies that bound to the RBD in convalescent sera did not target the ACE2IS (Fig. 2A and B). These data suggest that the ACE2IS is not a highly immunogenic epitope within the RBD.

**Figure 2.**
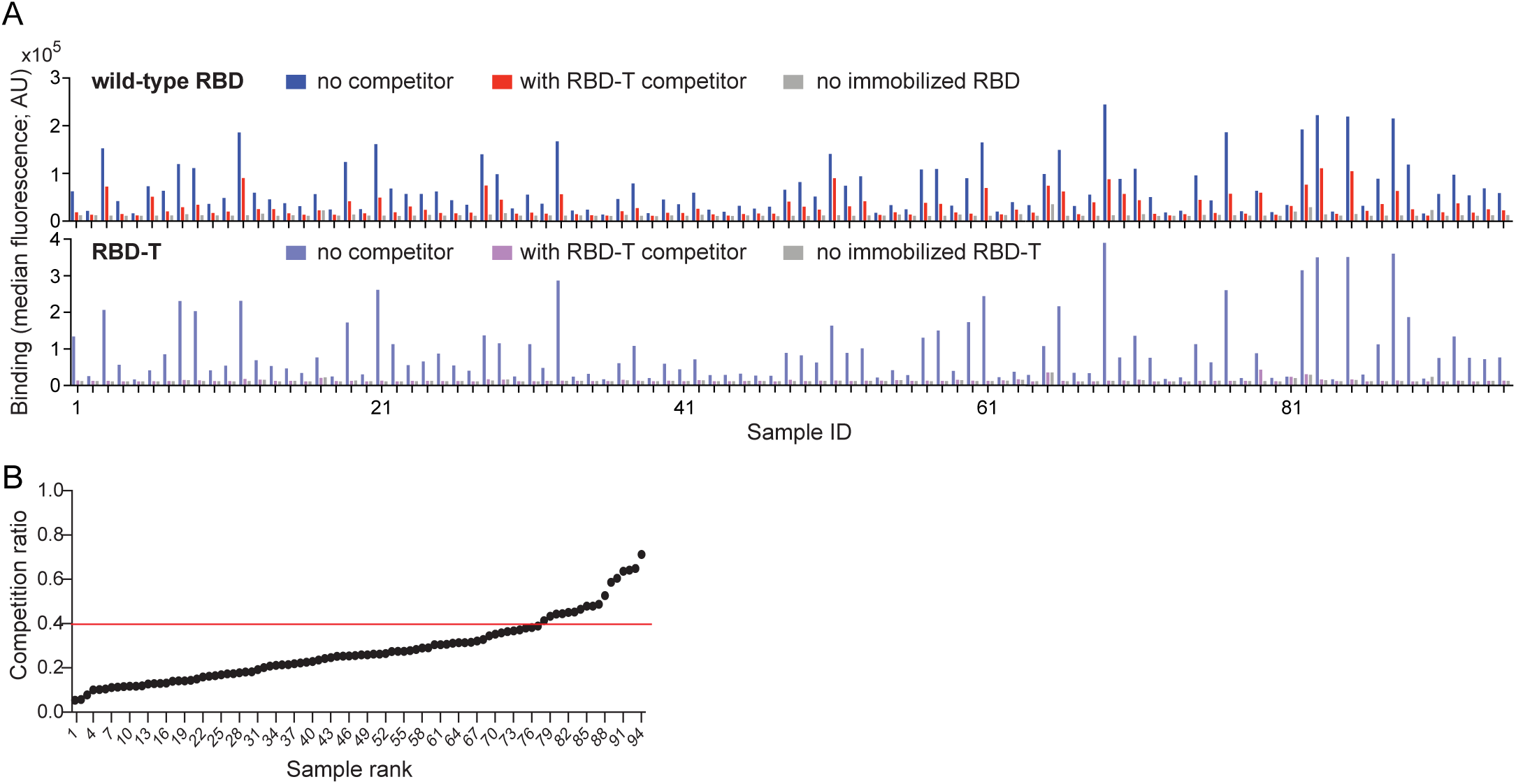
Majority of convalescent serum samples do not target the ACE2IS. (A) Binding analysis of IgGs in convalescent serum samples. In the top panel, the binding of the antibodies to the RBD in the absence (blue bars) and presence of soluble RBD-T as a competitor (red bars) are shown. The bottom panel shows a control experiment to confirm that competition with RBD-T was completely achieved in this assay. (B) The fraction of the antibodies targeting the ACE2IS among the RBD-binding antibodies in each serum sample is calculated as a competition ratio (signal in the presence of the RBD-T competitor over signal in its absence) and shown. Samples are sorted from low to high competition ratio. The red line indicates a competition ratio of 40%, i.e., 60% of binding signal is lost in the presence of the RBD-T competitor.

### Unbiased antibody selection for the spike protein and the RBD

To investigate the immunogenicity landscape of the spike protein and the RBD, we performed *in vitro* selection of a human antibody phage-display library against the spike protein and the RBD. The recovery of antibody clones in the *in vitro* selection primarily depends on the strength of binding to the antigen, and thus analyzing specificity and epitopes of enriched antibodies enables us to determine the immunogenicity landscape of the antigens from the point of the direct antibody-antigen interaction under well-define conditions. We used a synthetic antibody library that is constructed based on both intrinsic amino acid bias in immune repertoires and knowledge of structures and functions of naturally occurring antibodies [17, 18]. This, and closely related antibody libraries, have generated numerous high-affinity antibodies to diverse proteins [19].

We first performed selection for antibodies against the spike protein and the RBD in an unbiased manner, that is, we did not incorporate steps that might bias the recovery of antibodies binding to different epitopes (Fig. 3A). Clones from the 3rd and 4th rounds of selection were randomly picked, and their sequences and binding specificity were characterized in the phage-display format. We identified 95 unique clones out of 120 randomly picked colonies from the spike selection campaign (Fig. 3A). Strikingly, 89% of the clones bound to the spike protein but not the RBD or RBD-T (“Spike only” binders), indicating that these antibodies recognize the surfaces of the spike protein outside the RBD or the surfaces that partially consists of the RBD (Fig. 3A and B). Only 11% of the clones showed ability to bind to the spike protein as well as the RBD and RBD-T (“Spike+RBD” binders), and antibodies binding to the ACE2IS, as judged by the ability to bind to the RBD but not to RBD-T, were not identified from this selection. These results suggest that the RBD is not a highly immunogenic region of the spike protein.

**Figure 3.**
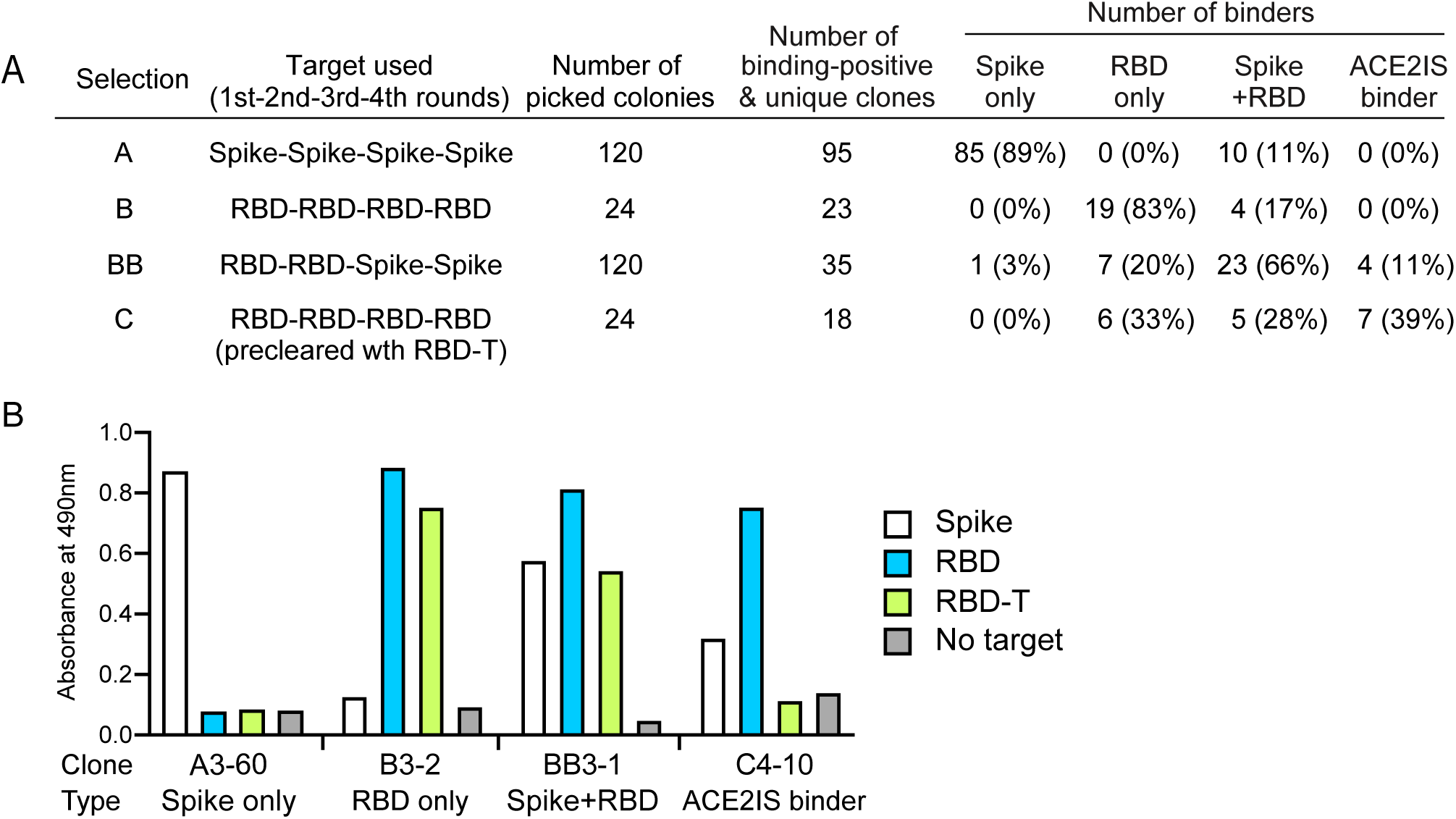
Antibody selections against the RBD and the spike protein revealed immunogenic properties of epitopes within the spike protein. (A) Summary of antibody selections in the biased and unbased manners. (B) Binding analysis of phage-displayed antibodies by ELISA. Data for representative clones for each specificity type are shown. See also Supplementary Figure 2 for the complete dataset.

From the RBD selection campaign, we identified 23 unique clones out of 24 randomly picked colonies (Fig. 3A). Although the ACE2IS is highly exposed on the RBD, antibodies targeting the ACE2IS were not identified from this selection. Interestingly, 83% of the clones showed binding to the RBD and RBD-T, but not to the spike protein (“RBD only” binders) (Fig. 3A and B). These data indicate that their epitopes are exposed when the RBD exists in isolation or after dissociation of the S1 subunit from the spike protein, but are not exposed when the RBD is a part of the fully assembled spike trimer. These results suggest that, if the RBD is used as an antigen for vaccination, antibodies targeting those epitopes of the RBD that are not exposed in the native spike protein can dominate in the initial immune response. Together, our data suggest that isolated RBD contains highly immunogenic epitopes that are inaccessible in the context of the fully assembled spike protein.

### Biased antibody selection readily identifies ACE2IS-targeting antibodies

Because the unbiased RBD selection campaign enriched antibodies that do not bind to the spike protein, we next performed antibody selection using both RBD and spike protein as the antigens (Fig. 3A) to intentionally enrich for clones that cross-react with both antigens. We identified 35 unique antibodies among 120 randomly chosen clones. A majority of clones bound to both RBD and spike protein (“Spike + RBD” binders), indicating that RBD contains epitopes that can be readily recognized in the context of the fully assembled spike protein (Fig. 3A and B). We note that antibodies that bound outside the RBD (“Spike only” binders) and to epitopes on the RBD that are inaccessible in the spike protein (“RBD only” binders) were still recovered, despite the use of a strong selection bias against such clones. We identified clones that bound to both RBD and spike protein but not RBD-T, indicating that they recognize the ACE2IS (“ACE2IS binders”). Interestingly, among RBD-binding antibodies, only 12% of clones recognize the ACE2IS in the RBD, whereas 88% of clones recognize epitopes other than the ACE2IS (Fig. 3A and B). These data are consistent with what we observed in the analysis of antibodies in convalescent sera (Fig. 2A and B) and suggest that the ACE2IS deflects antibody recognition.

We next performed selection of antibodies binding to the RBD that incorporated a negative selection step using RBD-T so as to enrich antibodies binding to the ACE2IS. From this selection, 39% of clones recognized the ACE2IS in the RBD, which is a substantially higher percentage than those recovered in the unbiased selections (0% for both selections using the spike protein and the RBD) as well as the RBD-spike selection (11%) (Fig. 3A). These data indicate that our phage-display library contains antibodies targeting the ACE2IS and such antibodies can be identified if we intentionally try to enrich such antibodies. Taken together, our data demonstrate that the RBD is not a dominant immunogenic domain in the spike protein and the ACE2IS is a minimally immunogenic epitope within the RBD.

Why is the RBD difficult to target when whole spike trimer is used as the antigen? Our data suggest that the spike protein has highly immunogenic surfaces outside the RBD, which enrich antibodies that target these epitopes and deplete antibodies to the RBD. The structure and conformational dynamics of the spike protein offer possible mechanistic basis for the low immunogenicity of the RBD. The RBD in the up conformation is largely exposed on the spike surfaces, whereas the RBD in the down conformation is buried in the spike protein (Fig.1A and 4A). Structural studies of the spike protein revealed that zero, one or two RBDs but not all three RBDs in the spike trimer form the up conformation at a given time [1, 2, 20, 21]. Consequently, the RBD is largely in the down conformation and most of the RBD is less exposed than the rest of the spike surfaces (Fig 4A). Furthermore, it has been reported that the RBD in the up conformation shows higher mobility than that in the down conformation [20]. Thus, both a lack of the complete exposure of the RBD and conformational mobility of the RBD are probably contributing to the challenges in recognizing the RBD by antibodies when the spike protein is used as the antigen.

**Figure 4.**
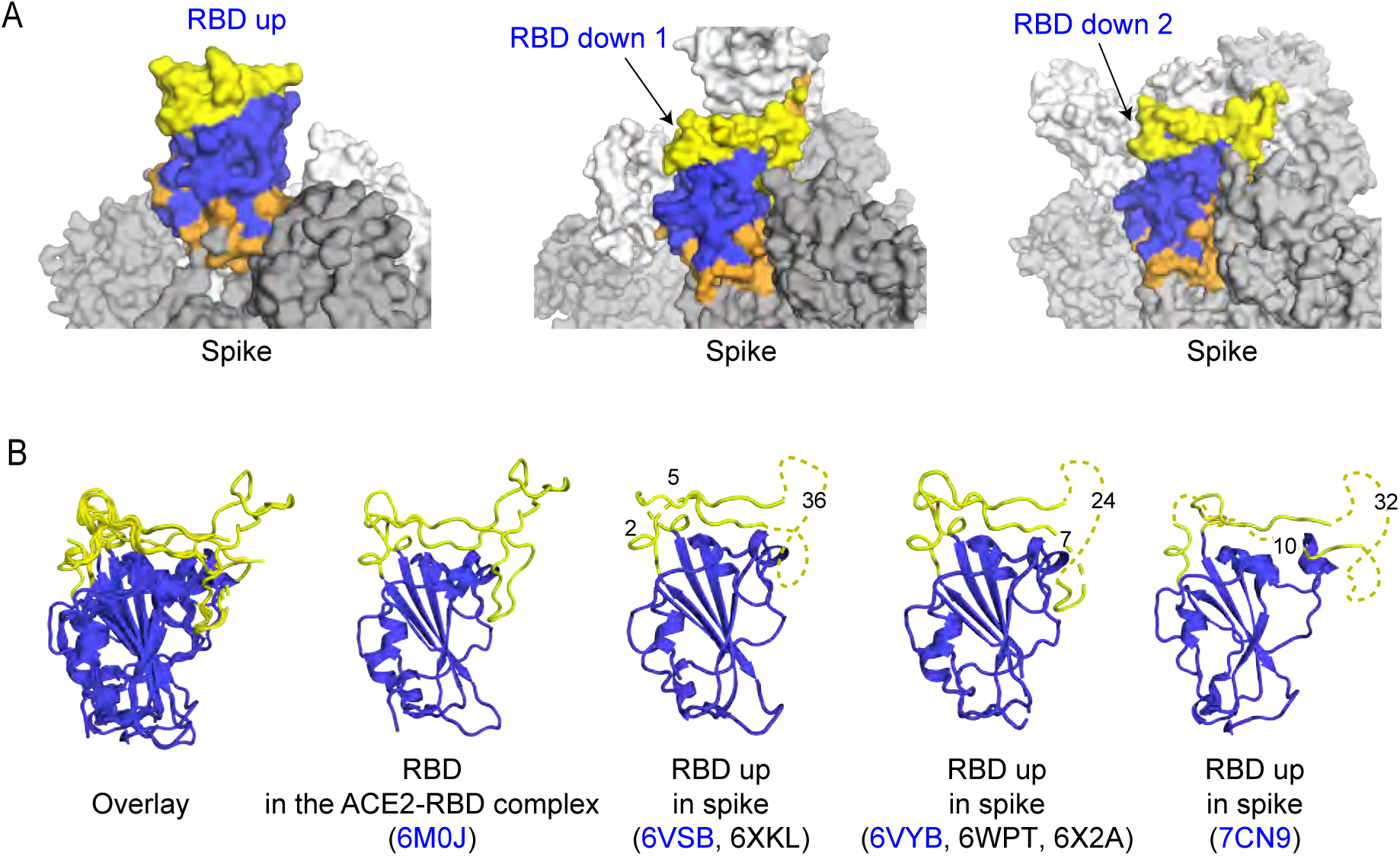
Possible mechanistic basis of low immunogenicity of the RBD and the ACE2IS. (A) The RBD in the up and the down conformations in the spike structure (PDB 6VSB). The RBD core and the receptor binding motif (RBM) are shown in blue and yellow. The residues within 6 Å of other regions in the spike protein are shown in orange. The other RBDs in the spike trimer are shown in white. The RBD in the up conformation is exposed whereas the RBDs in the down conformation are buried in the spike protein. (B) Comparison of the RBD structures. The RBD core and the RBM are shown in blue and yellow. The disordered regions in these structures are shown as the dotted lines, and the numbers of missing residues are indicated. The RBD structures indicated in a parenthesis have nearly identical structures, and the representative structures are shown and whose PDB codes are indicated in blue.

Our data indicate that the ACE2IS is minimally immunogenic within the RBD, although the ACE2IS is highly accessible when the RBD is in the up conformation. One possible explanation for the minimal immunogenicity of the ACE2IS is the conformational flexibility. Although the ACE2IS is well defined when bound to ACE2 or to an antibody, it is disordered in the free spike structures (Fig. 4B) [1, 2, 4, 22–25], suggesting that the ACE2IS is inherently flexible. It is difficult to generate antibodies to a flexible region, due to a large loss of conformational entropy upon binding that disfavors binding. Taken together, we propose that limited surface exposure and flexibility of the ACE2IS reduces its immunogenicity in the spike protein.

Our studies have important implications for vaccine design. Most of vaccine candidates utilize spike or the RBD as antigens (https://www.who.int/publications/m/item/draft-landscape-of-covid-19-candidate-vaccines). In our studies, 89% of antibodies from the unbiased selection for the spike protein recognize regions outside the RBD, and 83% of antibodies from the unbiased selection for the RBD recognize the surfaces on the RBD that are inaccessible in the context of the spike protein. These data imply that potent neutralizing antibodies targeting the ACE2IS may not be induced efficiently by only immunizing with the spike protein or only with the RBD. Consistent with this view, it has been reported that the majority of spike-binding antibodies that were generated in a patient during the first weeks of COVID-19 infection were not neutralizing and bind outside the RBD [26]. Similarly, Juno et al. observed a high frequency of B cells that target the spike protein outside the RBD in recovered patients with COVID-19 [15]. Our data demonstrate that the ACE2IS can be readily targeted by antibodies in the biased selection. Although such control of the human immune responses using designed antigens is impossible in the course of natural infection, we envision that a two-step vaccination scheme using the RBD followed by the spike protein could improve the efficiency of eliciting antibodies targeting the ACE2IS that confer strong viral neutralization.

## Materials and methods

### Plasmid construction

The amino acid sequence of the ectodomain of the spike protein (resides 16-1213) was collected from GenBank entry MN908947.3. A codon-optimized gene encoding the spike protein with modifications reported by Amanat et al [27], including stabilized mutations (K986P and V987P), the removal of the furin site (RRAR to A) and the addition of a T4 foldon trimerization domain, a hexahistidine tag and an Avitag at the C-terminus was synthesized (Integrated DNA Technologies) and cloned into the mammalian expression vector pBCAG. The codon-optimized genes encoding the receptor binding domain (RBD) of SARS-CoV-2 (residues 328-531; GenBank entry: MN908947.3) and a triple mutant (N487K/Q493K/N501K) of the RBD of SARS-CoV-2 with the hexahistidine tag and the Avitag at the C-terminus were synthesized (Integrated DNA Technologies) and cloned into the pBCAG vector.

### Mammalian cell culture

The A549 cells stably expressing the angiotensin-converting enzyme 2 (ACE2) on the cell surface were maintained in Dulbecco’s Modified Eagle’s Medium (DMEM, Thermo Fisher) supplemented with 2 μg/ml puromycin (InvivoGen), 10% fetal bovine serum (FBS, Gemini Bio) and penicillin/streptomycin (Thermo Fisher) at 37 °C with 5% CO_2_. The Expi293F cells (Thermo Fisher) were maintained in Expi293 Expression Medium (Thermo Fisher) at 37 °C with 8% CO_2_.

### Serum samples

Serum samples were collected from 94 Healthcare workers from NYU Langone Health who showed PCR-positive for COVID-19, as reported previously [14]. All patients gave written consent for this study and all samples were de-identified by following IRB #i20-00595.

### Expression and purification of antigens

The expi293F cells (Thermo Fisher) were transiently transfected with the vectors using the ExpiFectamine 293 Transfection kit (Thermo Fisher) according to the manufacturer’s protocol. The cells were incubated at 37 °C with 8% CO_2_ for 7 days. The cell culture supernatants were collected by centrifuge, and supplemented with a protease inhibitor cocktail (Roche) and 1mM PMSF. The recombinant proteins were purified from the filtrated supernatants using a HisTrap excel column (GE Healthcare). The purified proteins were biotinylated *in vitro* using the BirA enzyme in the presence of 0.5 mM Biotin and 10 mM ATP. The biotinylated proteins were further purified using the HisTrap excel column, and dialyzed into PBS. Purity of proteins were confirmed by SDS-PAGE analysis. The size exclusion chromatography using a Superdex 200 10/300 (GE Healthcare) for the spike protein and a Superdex 75 10/300 (GE Healthcare) for the RBD and the RBD triple mutant showed a monodispersed and single peak with the expected molecular mass (Supplementary Figure 1).

### Establishment of ACE2-expressing A549 cells

A549 cells were transfected with pLenti.ACE2 lentiviral vector that encodes ACE2 and puromycin resistance (Tada *et al*., Submitted). After two days, the cells were selected in medium containing 2 μg/ml puromycin and cloned at limiting dilution. Individual cell clones were screened by flow cytometry for high ACE2 expression and a single cell clone was expanded.

### Cell-based binding assay

The ACE2-A549 cells were detached from the flask using the citric saline buffer containing 15 mM sodium citrate and 135 mM KCl. The cells were washed with PBS twice and suspended in PBS-BSA (PBS containing 1% (w/v) BSA (Gemini Bio)). The cells were aliquoted into the wells of the 96-well round bottom plate (Greiner). The RBD and the RBD triple mutant were added to the wells containing the ACE2-A549 cells and incubated for 30min at room temperature. The cells were washed three times with 200 μL of PBS-BSA, and incubated with Streptavidin-Dylight650 (Thermo Fisher) for 30min at 4 °C. The cells were further washed three times with PBS-BSA and analyzed on an iQue screener (Satorius).

### Serum characterization

Antibody binding was characterized using a bead-based binding assay, generally following our previous publications [28, 29]. Biotinylated RBD was conjugated to Dynabeads M-280 streptavidin (Thermo Fisher Scientific) as follows. Ten microliters of the stock bead suspension were mixed with 90 µl of PBSB (PBS containing 0.1% (w/v) BSA (GeminiBio). The supernatant was removed using a magnetic stand and the beads were resuspended in 100 µl of PBSB. The beads were mixed with the equal volume of PBSB containing 15 nM of biotinylated RBD or biotinylated RBD-T and incubated at room temperature for 30 min. Biotin at a final concentration of 2.5 µM was then added to the solution and incubated for 5 min. The beads were washed once using a magnetic stand and resuspended at 0.5 mg/ml.

Five microliters of the beads were aliquoted in each well of a 96-well filter plate (Millipore), the liquid was removed using a vacuum chamber and the wells were washed once with 150 µl of PBST containing 1% (w/v) skim milk. Serum samples were heat-treated at 56°C for 1 hr and diluted 158-fold in PBST containing 1% (w/v) skim milk, and 25 µl each was added to the plate. For competition reactions, biotinylated RBD-T and streptavidin were mixed in such a way that all the biotin binding sites of streptavidin (tetramer) were occupied with RBD-T, and the complex was added to the serum solutions at a final concentration of 1 µM (calculated as monomer). After incubation for 30 minutes at room temperature with mixing on a shaker, the wells were washed twice with 150 µl of PBST-BSA (PBS containing 0.5% (w/v) BSA and 0.05% (v/v) Tween 20). The staining with a secondary antibody (anti-human IgG (FC gamma specific)-Alexa488, Jackson Immunoresearch), diluted 1/800 in PBS containing 1 % BSA, was performed in a total volume of 25 µl for 30 minutes at room temperature with shaking. The wells were washed again with PBST-BSA and the beads resuspended in 90 µl with PBST-BSA and analyzed on an iQue screener (Sartorius). Signals reported are the median fluorescence intensities.

### Phage display antibody selection

Sorting of a synthetic antibody library was performed as described previously [18].

Briefly, a synthetic human antibody library was sorted against the antigens listed in the Fig.3A. at the concentrations of 100 nM (1st round), 100 nM (2nd round), 50 nM (3rd round) and 20 nM (4th round). In the biased selection using RBD-T as the antigen for negative selection, biotinylated RBD-T was immobilized on the Streptavidin MagneSphere particles (Promega) and added to the solution containing phage. Phage bound to the RBD-T-beads were removed prior to the selection against the antigens.

### Phage ELISA

Phage ELISA was performed as described previously except that the 384-well plate was used instead of the 96-well plate [18]. Briefly, the wells of the 384-well ELISA plate (Nunc) were coated with neutravidin (Thermo Fisher) for 1 hr at room temperature. The wells were washed with PBS-T (PBS containing 0.1% Tween 20) and blocked with PBS containing 0.5% (w/v) BSA (Gemini Bio) for 1hr. After removing the blocking buffer, biotinylated antigens were added to each well and washed three times with PBS-T. The 5-fold dilution of the cell culture supernatants containing phage were added to the wells and incubated for 30 min. After washing the wells with PBS-T three times, anti-M13HRP (Sino Biological) was added to the wells. SIGMAFAST™ OPD (Sigma) was used as a substrate and 2 M HCl was used as a quenching solution. The absorbance at 490 nm was measured using a BioTek Epoch 2 plate reader (BioTek).

## Acknowledgments

We would like to thank all the volunteers for participation and Drs. Maria E. Aguero-Rosenfeld and Joan Cangiarella and NYU Langone Hospital Outpatient Laboratory for serum sample collection; Dr. Florian Krammer for making the preprint of his paper available; Dr. Jonathan Lai for guidance for the preparation of the spike protein. This work was supported in part by the National Institute of Health grants R21 AI158997, R01 CA194864 and R01 CA212608 to S.K., DP1 DA046100, R33 AI122390 and K08 AI120898 to N.R.L., and R21 CA246457 to T.H. K.W.T. was supported by the American Cancer Society fellowship PF-18-180-01-TBE.

## Notes

### Competing Interest Statement

The authors have declared no competing interest.

